# Neutral tumor evolution?

**DOI:** 10.1101/158006

**Authors:** Maxime Tarabichi, Iñigo Martincorena, Moritz Gerstung, Armand M. Leroi, Florian Markowetz, Paul T. Spellman, Quaid D. Morris, Ole Christian Lingjærde, David C. Wedge, Peter Van Loo, on behalf of the PCAWG Evolution and Heterogeneity Working Group

**Author notes:** To whom correspondence may be addressed: The Francis Crick Institute, 1 Midland Road, London, NW1 1AT, United Kingdom. These authors contributed equally. These authors jointly directed the work.

## Abstract

Tumor growth is an evolutionary process governed by somatic mutation, clonal selection and random genetic drift, constrained by the co-evolution of the microenvironment^1,2^. Tumor subclones are subpopulations of tumor cells with a common set of mutations resulting from the expansion of a single cell during tumor development, and have been observed in a significant fraction of cancers and across multiple cancer types^3^. Peter Nowell proposed that tumors evolve through sequential genetic events^4^, whereby one cell acquires a selective advantage so that its lineage becomes predominant. According to this traditional model, the selective advantage is conferred by a small set of driver mutations, but, as the subclones that bear them expand successively, they accumulate passenger mutations as well, which can be detected in sequencing experiments^1^. Genomes of individual tumors contain hundreds to many thousands of these genetic variants, at a wide range of frequencies^5,6^. Given that genetic drift alone can drive novel variants to high frequencies, it is of great interest to discern the relative importance of selection and drift in shaping the frequency distribution of variants in any given tumor.

Williams *et al*.^7^ recently proposed a way to do so. They found that a simple model of tumor growth in which all novel variants are selectively neutral, that is, whose dynamics are governed entirely by drift, predicts a linear relationship between the number of mutations *M*(*f*) present in a fraction *f* of cells and the reciprocal of that fraction: 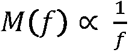. They argued that deviation from this null model, i.e. the R-squared of the linear fit is below the minimum observed in neutral simulations (R^2^ < 0.98), indicates the presence of selection and that this can be tested by means of variant allele frequencies (VAFs) from which *f* can be derived. Applying this rationale to real cancer data from The Cancer Genome Atlas (TCGA), the test proposed by Williams *et al*. did not reject the null model, that is neutrality, in about one third of the cases and the authors concluded that these tumors are neutrally evolving. More recently, multiple myelomas with evidence for the proposed linear relationship were associated with poorer prognosis^8^.

While providing an interesting approach to infer selection in human cancers, unfortunately four major simplifying assumptions underlie the analysis by Williams *et al*. that might render the conclusions questionable.

First, inferring *f* of variants from their VAF requires accurate estimates of local copy number, overall tumor purity and ploidy. Williams *et al*. attempted to account for some of these factors by restricting their analyses to variants with VAF between 0.12 and 0.24 and located in copy-neutral regions of the genome. However, even in that limited VAF window, the VAF of a mutation does not reflect its true *f* in many cases. For example, in tumors with whole genome duplications, i.e. 37% of tumors in the analyzed dataset^9^, the peak of clonal mutations acquired after the whole genome doubling event is at or below VAF = 0.25 (one out of four copies in a 100% pure tumor sample), which would lead to artificial deviation from the linear fit within that VAF window.

Second, the interpretation of the analyses is inconsistent with the use of neutrality as a null model. Failure to reject the null hypothesis is not the same as proving it true, i.e. that all neutral simulations have R^2^ > 0.98 does not prove that non-neutral simulations would never yield R^2^ > 0.98. One would need to demonstrate that this condition is sufficient to infer neutrality but also, no equally suited models of non-neutral tumor growth should yield R^2^ > 0.98.

To assess this, we simulated simple tumor growth in which we explicitly model one subclonal expansion with a selective advantage, i.e. increasing its division rate *λ* and/or the mutation rate *μ* of the subclone (**Supplementary Methods**). Using the original method described by Williams *et al*., neutrality is rejected only within a narrow range of *λ* and *μ* values tested that would lead to detectable subclones (true rejection of neutrality in ~11% of simulations; **Fig. 1a**). We conclude that a linear fit with R^2^ > 0.98 is not sufficient to call neutrality and that improper use of this model could result in substantial over-calling of neutrality.

**Figure 1.**
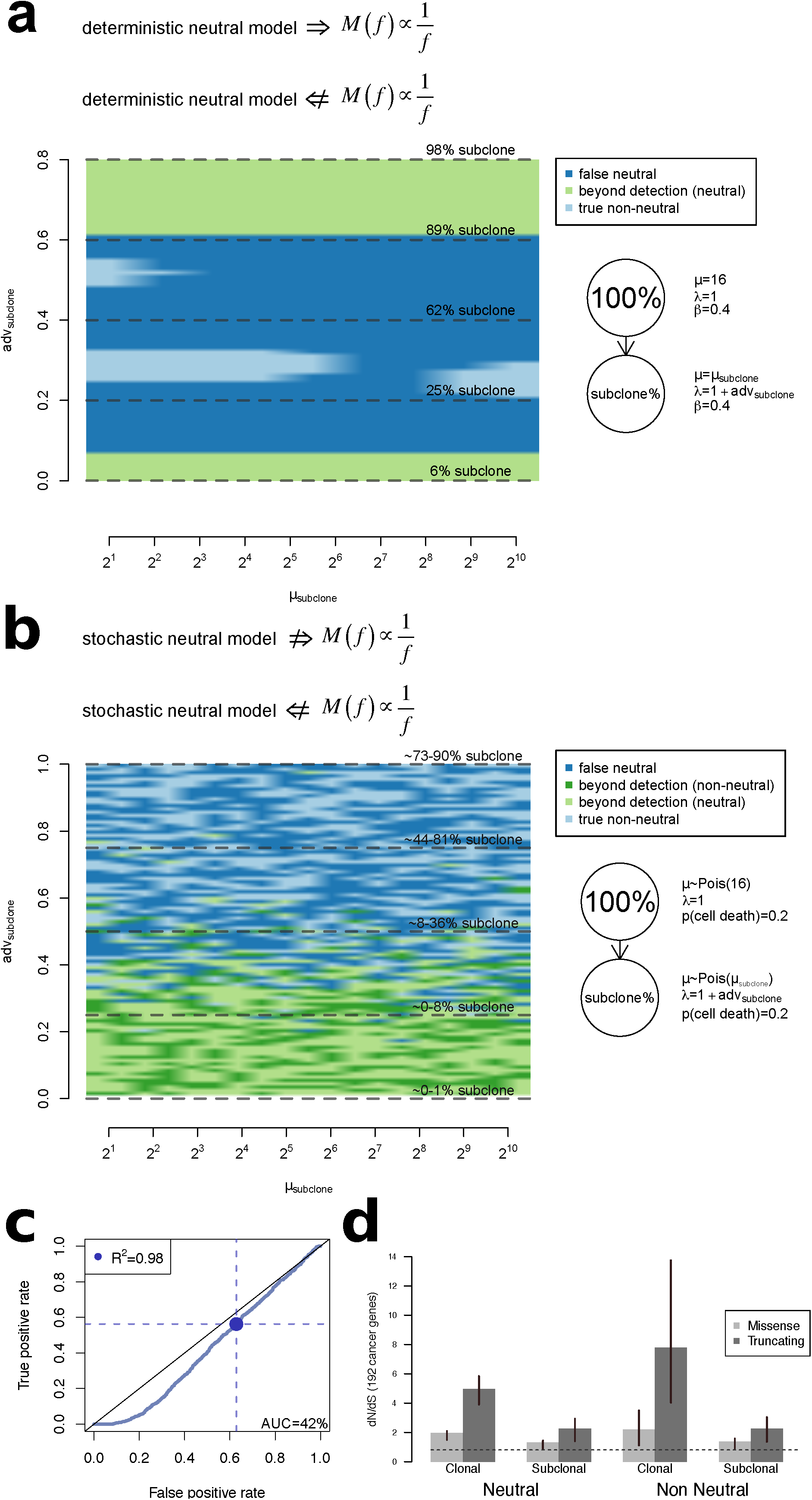
**(a) Neutrality calls in simulations of tumor growth with subclonal expansion underlying selective sweeps**. The tree topology being modelled is represented on the right together with the parameters of the neutral evolution equations for the two subpopulations of cells (**Supplementary Methods**). The subclone’s fraction (subclone %) increases with its selective advantage adv_subclone_. We vary the *λ* = 1 + adv_subclone_ and *μ* parameters of the subclone along a grid. Simulations are defined as true non-neutral (light blue) or false neutral (dark blue) when the growing subclone has expanded sufficiently to be detectable and the sweep is not complete, i.e. 10% ≤ subclone % ≤ 90%, otherwise the subclone is considered beyond detection (light green). Non-neutral call: R^2^ < 0.98; neutral call: R^2^ ≥ 0.98**. (b) As (a), using the Gillespie algorithm to simulate branching processes**^10^. Simulations leading to subclones beyond detection are either called neutral (light green) or non-neutral (dark green). Because of the stochastic nature of branching processes, different subclone % values are obtained across simulations from the same adv_subclone_ values. For five increasing adv_subclone_ values, we report median ± mad of the subclone % across the simulations. **(c) Summary ROC curve for the neutral vs. non-neutral classification based on the R^2^ values in 1,919 non-neutral simulations from (b), and 1,919 simulations of neutral tumors**. The false positive rate and the true positive rate are highlighted for R^2^ = 0.98 used by Williams *et al*. **(d) dN/dS analysis**. Maximum likelihood estimates of the dN/dS ratios and associated 95% confidence intervals for (sub)clonal mutations in TCGA tumors categorized into neutral and non-neutral groups. Ratios for missense and truncating mutations are given. dN/dS > 1 indicates positive selection.

Third, the deterministic model of tumor growth described by Williams *et al*. relies on strong biological assumptions, among which are synchronous cell divisions, constant cell death and constant mutation and division rates. Stochastic models of tumor growth are biologically more realistic, as they allow for asynchronous divisions and probabilistic mutation acquisition, cell death and division rates. Using simple branching processes to simulate neutral and non-neutral growth^10^ (**Supplementary Methods**), we show that R^2^ > 0.98 for 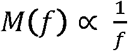 is neither a necessary nor a sufficient property of neutrally evolving tumors (**Fig. 1b**). Although it can be shown that the expected cumulative number of mutations - i.e. the average over many independent samples 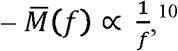 due to the biological noise modeled in branching processes, a typical realization of the neutral process in a single sample deviates substantially from the expected linear fit, rendering an R-squared threshold inaccurate to infer neutrality. As a result, discrimination of neutral and non-neutral simulated tumors using a linear fit is almost arbitrary, with 53.5% false positive neutral calls in non-neutral tumors (**Fig. 1b**) and an area under the ROC curve of 0.42 for the classification of 1,919 neutral and 1,919 non-neutral tumors (**Fig. 1c**).

Fourth, we reason that in tumors called neutral, no subclonal selection should be detected. To evaluate this, we use an orthogonal method to identify selection, based on the observed variants themselves rather than on their allele frequencies. dN/dS analysis derives the fraction of mutated non-synonymous positions to the fraction of mutated synonymous positions in the coding regions. It has been widely used to detect the presence of negative or positive selection of non-synonymous variants in coding regions^11,12^. We applied a dN/dS model optimized for the detection of selection in somatic cancer variants^13^ to TCGA exome data using a published list of 192 cancer genes^14^ (**Supplementary Methods**). The analysis was performed separately using variants called as clonal or subclonal (**Supplementary Methods**), in tumors called neutral and non-neutral based on the rationale outlined by Williams and colleagues^7^. dN/dS ratio analysis revealed significant positive selection in subclonal mutations of tumors classified as neutral (**Fig. 1d**), further suggesting that the approach described by Williams *et al*. is under-equipped to detect the presence or absence of selection.

In summary, Williams *et al*. proposed that about one third of tumors are neutrally evolving. However, we highlight four simplifying assumptions - to our knowledge not previously highlighted - and find that the proposed approach will often identify individual tumors as neutral when they are non-neutral and non-neutral when they are neutral. A new paper by the same group^15^ introduces a Bayesian test for detecting selection from VAFs. The test estimates selection coefficients and, as such, is an important advance over Williams *et* al.’s frequentist test, which does not. The authors acknowledge that the test can only detect large fitness differences, but nevertheless call tumors that fail it “neutral’’ when they are merely those in which a weak test has failed to detect selection. We note that neutral theory has been developed in population genetics, ecology and cultural evolution and that similar tests have been proposed in all of these fields and, in all, eventually been found wanting for the same reason: variant abundance distributions do not contain enough information to exclude selection^16-18^. It is of clinical importance to identify and better understand the drivers of the potentially more aggressive (sub)clones expanding under selective biological or therapeutic pressure, as these are good candidates for predicting resistance and exploring combination therapy. Williams *et al*. are to be commended for having introduced explicit neutral tumor growth models into tumor genomics. However, quantifying the relative importance of drift and selection in shaping the allele frequencies of single tumors clearly remains an open challenge. Studies relying on their proposed test (e.g. ^8^) might, then, need reevaluation.

## Acknowledgments

This work was supported by the Francis Crick Institute, which receives its core funding from Cancer Research UK (FC001202), the UK Medical Research Council (FC001202), and the Wellcome Trust (FC001202). MT is a postdoctoral fellow supported by the European Union’s Horizon 2020 research and innovation program (Marie Skłodowska-Curie Grant Agreement No. 747852-SIOMICS). PVL is a Winton Group Leader in recognition of the Winton Charitable Foundation’s support towards the establishment of The Francis Crick Institute. IM is funded by a Cancer Research UK Career Development Fellowship (C57387/A21777). DCW is funded by the Li Ka Shing foundation. This work was supported by grant 1U24CA210957 to PTS. FM would like to acknowledge the support of The University of Cambridge, Cancer Research UK and Hutchison Whampoa Limited. Parts of this work was funded by CRUK core grant C14303/A17197. This project was enabled through access to the MRC eMedLab Medical Bioinformatics infrastructure, supported by the Medical Research Council (grant number MR/L016311/1). Parts of the results published here are based upon data generated by the TCGA Research Network: http://cancergenome.nih.gov/.

## Author contribution

MT, IM, MG, AML, FM, PTS, QDM, OCL, DCW, PVL participated in argumentation. MT, OCL, DCW and PVL derived the deterministic equations. MT wrote the code and generated the figures, with input from IM, MG, OCL, DCW and PVL. MT, OCL, DCW, PVL drafted the manuscript, revised by IM, MG, AML, FM, PTS, and QDM. All authors read and approved the manuscript.

## Methods

### Outline

First, we describe the two tumor growth models that were used. The first is based on the deterministic continuous model presented by Williams *et al*.^1^. The second is based on a branching process, a commonly used discrete and fully stochastic growth model. We next explain how, using these two models, we can simulate variant allele fractions encountered in tumor sequencing studies. We describe our implementation of the approach by Williams *et al*.^1^ to infer the most likely evolutionary path after the emergence of the most recent common ancestor (MRCA), i.e. neutral *vs*. non-neutral evolution. Finally, using real data from The Cancer Genome Atlas, we compare neutrality calls to results of dN/dS analysis, an independent and well-established approach to detect selection. We further describe the availability of the code as a tarball containing R and Java scripts and a Java runnable jar file called via one of the R scripts.

### Simulations - continuous deterministic models

The deterministic equations described in Williams *et al*.^1^ relate the number of cells in a tissue growing exponentially, *N*,

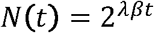

and the cumulative number of mutations, *M:*

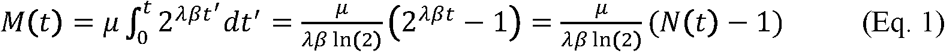

at any given time *t ≥ 0*, where *λ > 0* is the division rate per unit of time, *β ≥ 0* is the unitless “effective” division fraction, i.e. the fraction of divisions in which both daughter cells survive (β = 1 for no cell death, β < 1 to model cell death), and μ > 0 is the mutation rate per cell division.

We have used these continuous deterministic models to simulate tumor growth *in silico* and followed each mutation and its corresponding variant cell fraction. To derive the cell fractions, we follow the progeny of the mother cell within which each mutation occurred.

Assume that the MRCA appears at time *t_1_*, with division coefficient *β_1_*, division rate *λ_1_*, and mutation rate *μ_1_*. To model a selective sweep within the cell population spawned from the MRCA, we assume that at time *t_2_ > t_1_*, a subclone is initiated with division coefficient *β_2_* division rate *λ_2_*, and mutation rate *μ_2_*.

There is positive selection when *λ_2_β_2_* > *λ_1_β_1_*. At time *t* the number of cells spawned from the MRCA but not part of the subclone (i.e. the cells with parameters *β_1_, λ_1_, μ_1_;* further referred to as the MRCA lineage) is

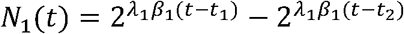

where the second term is omitted when *t < t_2_*. Similarly, the number of cells at time *t* from the subclonal lineage (i.e. with parameters *β_2_, λ_2_, μ_2_*) is

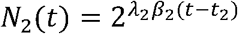

when *t > t_2_* and *N_2_*(*t*) *= 0* otherwise. The total cell count at time *t* is

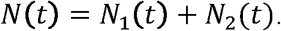

The tumor growth simulation is terminated at time *T* > *t_2_* and we derive the distribution at time *T* of the cell fractions for all mutations in the tumor.

### Following the number of mutations and their cell fraction

Because the equations are continuous, they can lead to non-integer numbers of mutations and cell divisions. Hence, rather than deriving the number of mutations and their allele frequencies *f* at discrete time points, we model divisions in continuous time. We assess the number of additional mutations that have been added in fixed (small) time intervals of length *dt*. From Eq. (1), we find that the number of additional mutations occurring in the time interval [*t*, *t* + *dt*] within a population of cells from the same lineage (i.e. parameters *β*, division rate *λ*, and mutation rate *μ*) is:

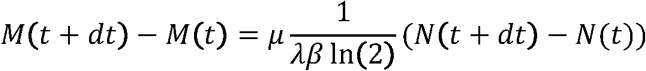

For a mutation occurring at time *t*, we may compute the variant cell fraction at time *T*. If the mutation occurred in a cell from the MRCA lineage that was not inherited by the subclone-initiating cell, then the variant cell fraction is

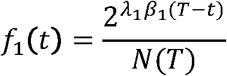

If the mutation occurred in the subclone, then the variant cell fraction is

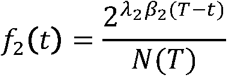

Finally, if the mutation occurred in an ancestor cell of the subclone-initiating cell, then the variant cell fraction is

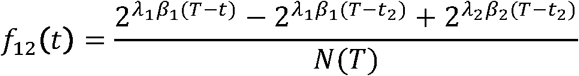

Alternatively, we may calculate variant cell fractions in two steps, first determining the variant cell fraction of a mutation within the subpopulation of cells from the same lineage, and then scaling the variant cell fraction by the size of that subpopulation relative to the total cell population.

### Setting the parameters for the grid of simulations

In each of our simulations the subclone growing under selective advantage appears at the 11^th^ generation and the tumor is sampled at the 40th generation with a virtual purity of 100%. The number of initial clonal mutations *μ_0_* is not part of these models, and we arbitrarily set *μ_0_* = *μ*_2_. We fix the following parameters: clonal mutation rate *μ_1_* = 16, clonal division rate *λ_1_* = 1, clonal division efficiency *β_1_* = 0.4, subclonal *β_2_* = 0.4. The depth of sequencing of the variants cov ~ Pois(10,000) to approach the theoretical distribution and the alternate read counts ~ Bin(cov, *f/2*), where *f* is the variant allele frequency derived from the model (see section on simulating tumor variant allele frequencies from sequencing data). We explore the results of the neutrality calls for a grid of parameter values wide enough to encompass many realistic combinations:

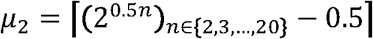

and

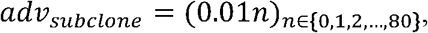

where

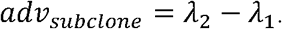

### Simulations – fully stochastic models

To model stochastic discrete tumor growth, we use branching processes with the Gillespie algorithm^2^. These simulated tumors grow under asynchronous division, with zero or one subclone.

This was coded in Java. Each cell is a Java object and has four attributes: a Boolean value reporting whether the cell is alive or dead; an integer for the average number of mutations per division; an integer with mother cell ID; and an *ArrayList* of all *MutationSets* inherited from the mother cell. *MutationSet* is another class, for which each object contains one integer for the mother cell ID and one integer for the number of mutations within them. The constructor of *MutationSet* takes the mutation rate of the mother cell as average number of events per interval of a Poisson distribution to draw the number of mutations.

Starting with an *ArrayList* of one tumor initiating cell, for each of 2^20^ cell division events, one cell is picked randomly from the living cells and either dies with probability P(*cell death*) or divides into two daughter cells with probability *P*(*division*) = 1 - P(*cell death*), akin to the Gillespie algorithm.

In our simulations, the subclone appears at the 2^8 th^ division (~8^th^ generation) by changing the division rate value of one of the cells, and the tumor is sampled at the 2^20 th^ division (~20^th^ generation). In these simulations, the number of mutations acquired at each cell division for each daughter cell is drawn from a Poisson distribution for the MRCA lineage *μ* ~ Pois(*μ*_MRCA_) and the subclone lineage *μ* ~ Pois(*μ*_subclone_).

The subclone is selected for division with probability

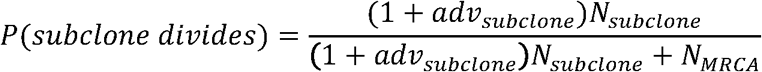

where *N_subclone_* and *N_MRCA_* are the number of cells from the subclonal lineage and the MRCA lineage, respectively, and *adv_subclone_* > 0 for positive selection and *adv_subclone_* = 0 for neutral growth. The MRCA population will be selected for division with probability 1 - P(*subclone divides*).

Within the selected clone, one cell is selected randomly for division with probability

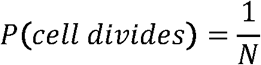

where *N* = *N*_MRCA_ if the cells belong to the MRCA lineage or *N* = *N_subclone_* if the cell belongs to the subclonal lineage.

With higher P(*cell death*), the first divisions are more likely to lead to the death of all cells and the tumor quickly stops growing. To limit this effect when cell death is high, we force the *D* first divisions to happen, i.e. P(*cell death*) = 0 transiently until at least *2D* cells are alive.

### Setting the parameters for the grid simulations

In our simulations, starting from one tumor initiating cell, for each of the 2^20^ cell division events, one cell is picked randomly and either dies with probability P(cell death) = 0.2 or divides into two daughter cells with probability P(division) = 1 - P(cell death) = 0.8. The subclone appears at the 2^8 th^ division (~8^th^ generation) and the tumor is sampled at the 2^20^-th division (~20^th^ generation). The ancestor clone’s mutation rate *μ* ~ Pois(16). The average depth of coverage is 100X (see section on simulating tumor variant allele frequencies from sequencing data). In our simulations, *D* = 6.

We explore a grid of values for

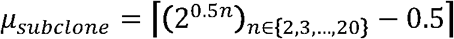

and

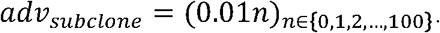

This leads to 19*101=1,919 simulated tumor simulations covering the grid.

### Simulating tumor variant allele frequencies from sequencing data

Using the tumor growth models presented here, we can derive the exact number of mutations and their prevalence within a virtual tumor. These are taken as input to simulate the frequencies that would be observed in the sequencing reads from real tumor tissue.

In order to test the initial hypothesis, i.e. 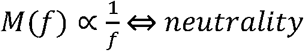, we start with the simplest models and assume: (i) the absence of non-tumor contaminant, (ii) 100% of the tumor cells are resected, and (iii) a fully diploid cancer genome.

Given exact cell fractions, *f* of each mutation and an average sequencing coverage, *cov*, we draw for each individual mutation the total number of reads covering its genomic position *N* from a Poisson distribution *N* ~ Pois(cov), and the alternate read counts *alt* ~ Bin(*N,f/2*), where *f/2* is the allelic fraction for diploid regions. Finally, we generate variant calls by taking mutations with *alt* > 2 and derive the variant allelic fraction (VAF) of each variant 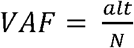. We then use the VAF distribution to call neutral and non-neutral tumors, as described by Williams *et a*l.^1^

### Calling neutral tumors

We followed the description by Williams *et al*.^1^ to call neutral and non-neutral tumors based on the variant allele frequencies of their somatic single nucleotide variants. Tumors with less than 12 mutations with 0.12 ≤ VAF ≤ 0.24 were removed. From the TCGA dataset, only tumors with a purity of at least 70%, as inferred by ASCAT^3^, were analyzed.

We calculated the explained variance (R^2^) for linear regression models both with fixed intercept (intercept = 0) and without fixing the intercept, using the R commands:

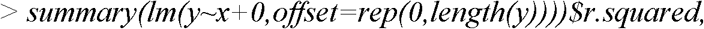

and

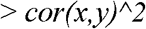

respectively, where *y* is the cumulative number of mutations and *x* is the inverse allelic frequency minus the upper limit 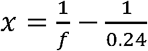. Results presented in the manuscript were obtained using a variable intercept. In **Supplementary Fig. 1**, we show the heat map of **Figure 1a** using a fixed intercept. Both methods show 97.5% agreement (**Supplementary Fig. 2**).

**Supplementary figure 1.**
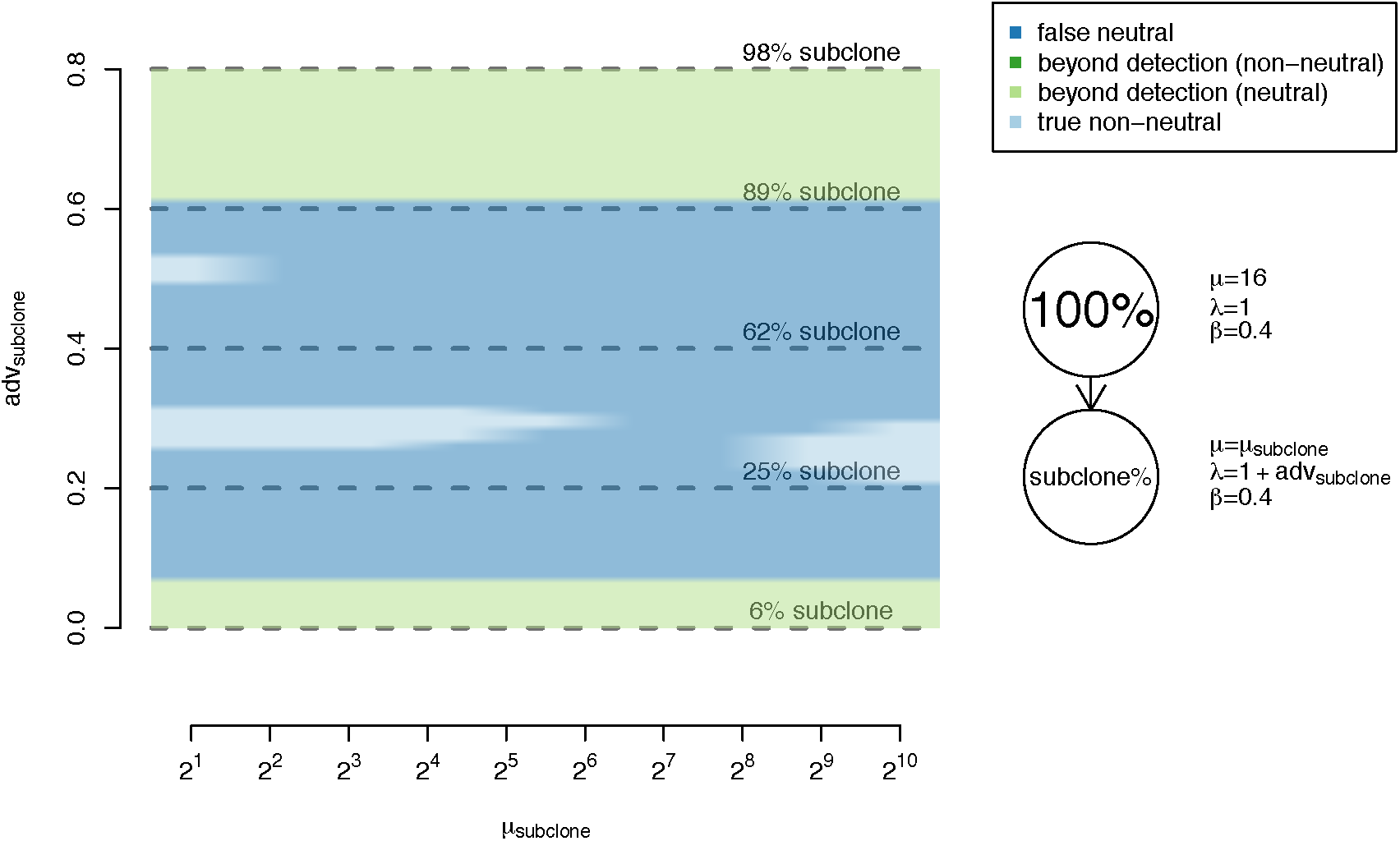
As reported in figure 1a using R^2^ of a linear regression with fixed intercept = 0. The tree topology being modelled is represented on the right together with the parameters of the neutral evolution equations for the two subpopulations of cells. The subclone’s fraction (subclone %) increases with its selective advantage adv_subclone_. We vary the *λ* = 1 + adv_subclone_ and *μ* parameters of the subclone along a grid. Simulations are defined as true non-neutral (light blue) or false neutral (dark blue) when the growing subclone is sizable enough to be detected and the sweep is not complete, i.e. 10% ≤subclone % ≤ 90%, otherwise the subclone is considered beyond detection (light green). Non-neutral call: R^2^ < 0.98; neutral call: R^2^ ≥ 0.98.

**Supplementary figure 2.**
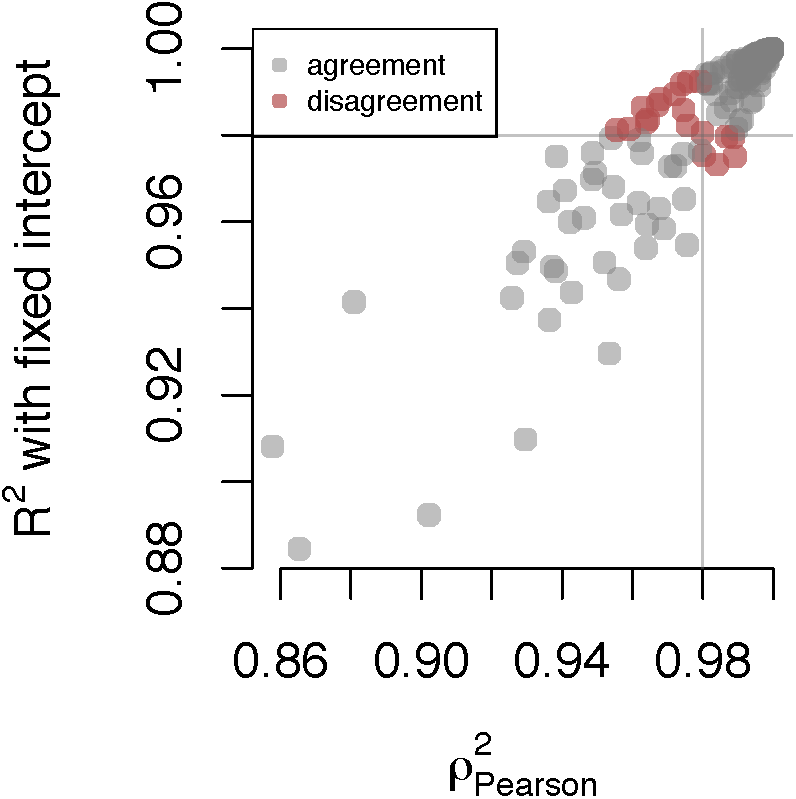
R^2^ values for the same simulations as in Supplementary figure 2, with variable and fixed intercept, showing an agreement of 97.5% on the neutral calls. The x-axis represents R^2^ values (squared Pearson’s correlation coefficients) for the linear regression between *M*(*f)* and *f* for the simulations in **Supp. Fig. 1**. The y-axis represents R^2^ values with fixed intercept = 0. Neutral calls, made if R^2^ ≥ 0.98, agree for 97.5% of these simulations (grey) and disagree for 2.5% of them (red).

### ROC and area under the curve

Using fully stochastic branching processes, we simulated 1,919 non-neutral tumors and 1,919 neutral tumors and derived the R^2^ values of the linear fit between the cumulative number of mutations and their inverse variant allelic fraction (VAF) within 0.12 ≤ VAF ≤ 0.24. We then plotted the ROC using the R package ROCR version 1.0–7 and calculated the false positive rate and the true positive rate assuming the R^2^ = 0.98 threshold used by Williams *et al*.^1^

### Detection of selection in neutral and non-neutral tumors - dN/dS

#### Dataset

We ran our analyses on the data from The Cancer Genome Atlas, using CaVeMan^4,5^ single nucleotide variant calls, and ASCAT^3^ copy number calls, as described by Martincorena *et al*.^6^

#### Grouping variants into clonal and subclonal categories

To classify variants as clonal or subclonal, we used a one-sided proportion test to assess whether the alternate and total read counts of each variant were compatible with its clonality, given its underlying number of DNA copies, and the overall tumor purity. This method is previously described in Alexandrov *et al.^7^*

#### dN/dS analysis and control gene sets

We performed dN/dS analysis to detect positive or negative selection of non-synonymous variants, as described by Martincorena *et al.^6^* The R package dNdScv was used to derive the dN/dS values and is available on github: https://github.com/im3sanger/dndscv. We ran dN/dS separately on clonal and subclonal mutations and separately in the neutral and non-neutral tumors, using a published list of 192 cancer genes (COSMIC v.80 - cancer.sanger.ac.uk^8^)^a^.

As a control, we ran dN/dS on subclonal mutations using 100 random sets of 192 genes, uniformly sampled from 20,090 annotated genes from hg19^6^. The 95% interval of dN/dS values was above 1, i.e. showed evidence for positive selection, in 3 out 100 random gene sets. We further reasoned that not all genes are equally “important” to the 192 COSMIC genes across tissues and took their gene expression across tissues as a proxy for their importance. We downloaded the human bodymap 2.0 (https://www.ebi.ac.uk/gxa/experiments/E-MTAB-513/Results) TPM matrix of expression and ranked genes by their values. We summed the ranks across relevant tissues (adrenal gland, brain, breast, colon, kidney, leukocyte, liver, lung, ovary, prostate gland, thyroid gland) and rerun dN/dS on 4×100 192-gene sets randomly sampled from the 10,000, 5,000, 2,500 and 1,000 top-ranked (highly-expressed) genes in those tissues. Among these four lists of highly expressed genes, the 95% intervals of dN/dS values were >1 in 2, 6, 4, 5 out of the 4×100 gene sets, respectively, confirming that the dN/dS signal is (cancer-)gene specific and is not biased in random gene sets. We then reasoned that gene co-expression levels might be a better proxy for “cancer-relevance” of the genes. To this end, the tool Gemma^9^ (https://gemma.msl.ubc.ca/home.html) was run to identify genes showing evidence for co-expression with one of the 192 cancer genes in >21 out of 442 gene expression datasets from the *Master set for human*. This identified 2,089 unique co-expressed genes (with a median of 2 co-expressed genes per cancer gene), from which we removed the 227 genes overlapping with the 719 cancer genes from the most recent cancer gene census (COSMIC v.84 - cancer.sanger.ac.uk^8^). We then sampled 192 unique genes from the 1,862 genes with probabilities of each gene *g* being sampled 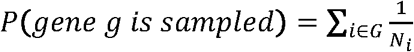, where G are the genes from the 192 genes that are co-expressed with *g*, and *N_i_* is the number of co-expressed genes with gene *i*. We ran dN/dS again on 100 of these 192-gene sets and found that 4 and 5 out of 100 gene lists yielded 95% confidence intervals of dN/dS >1 for subclonal mutations in neutral and non-neutral tumours, respectively.

### Effect of copy number

We repeated the analyses after selecting only variants that fall within diploid regions, i.e. 1 copy of allele A and 1 copy of allele B according to ASCAT^3^, to show that the results were not induced by unreliable neutral calls, which could have resulted from the distortion of allele frequencies by copy number changes (**Supplementary Fig. 3**).

**Supplementary figure 3.**
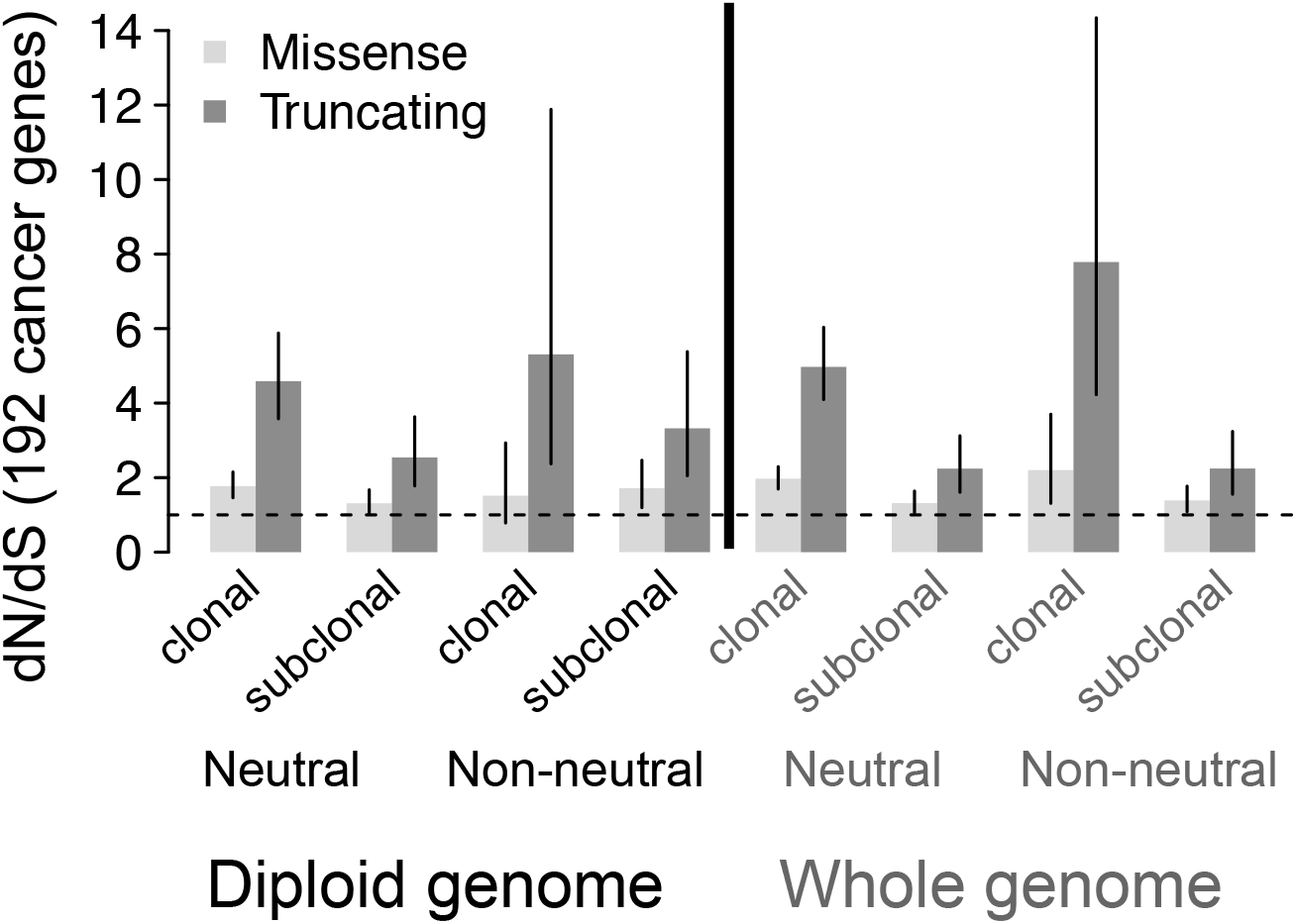
dN/dS ratios on all mutations vs. mutations in diploid regions only. Maximum likelihood estimates of the dN/dS ratios and associated 95% confidence intervals for (sub)clonal mutations in TCGA tumors categorized into neutral and non-neutral groups. Ratios for missense and truncating mutations are given. dN/dS > 1 indicates positive selection.

### Code reproducibility and availability

Analyses and figures were generated using R version 3.1.3. The branching processes are coded in Java. The code for simulations is available as a tarball (included within this submission) with R scripts for the deterministic simulations and for deriving the figures, and a Java runnable jar file for generating variant fractions from the branching processes together with the associated Java source code.

## Footnotes

a ABL1, ACVR1, ACVR1B, AKT1, ALK, AMER1, APC, AR, ARID1A, ARID2, ASXL1, ATM, ATP1A1, ATP2B3, ATR, ATRX, AXIN1, AXIN2, BAP1, BCOR, BIRC3, BRAF, BRCA1, BRCA2, CACNA1D, CALR, CARD11, CASP8, CBL, CBLB, CD79A, CD79B, CDC73, CDH1, CDKN2A, CDKN2C, CEBPA, CIC, CNOT3, COL2A1, CREBBP, CRLF2, CSF1R, CSF3R, CTNNA1, CTNNB1, CUX1, CXCR4, CYLD, DAXX, DICER1, DNM2, DNMT3A, EGFR, EML4, EP300, ERBB2, ERG, ESR1, ETNK1, EZH2, FAT1, FAT4, FBXO11, FBXW7, FGFR1, FGFR2, FGFR3, FLT3, FOXA1, FOXL2, FUBP1, GATA1, GATA2, GATA3, GNA11, GNAQ, GNAS, GRIN2A, H3F3A, H3F3B, HIF1A, HIST1H3B, HNF1A, HRAS, IDH1, IDH2, IKBKB, IKZF1, IL6ST, IL7R, JAK1, JAK2, JAK3, KCNJ5, KDM5C, KDM6A, KDR, KIT, KLF4, KMT2C, KMT2D, KRAS, MAP2K1, MAP2K2, MAP2K4, MAX, MED12, MEN1, MET, MLH1, MPL, MSH2, MSH6, MTOR, MYD88, MYOD1, NF1, NF2, NFE2L2, NFKBIE, NOTCH1, NOTCH2, NPM1, NRAS, NT5C2, NTRK3, PAX5, PBRM1, PDGFRA, PHF6, PHOX2B, PIK3CA, PIK3R1, PLCG1, POLE, POT1, PPP2R1A, PPP6C, PRDM1, PRKACA, PRKAR1A, PTCH1, PTEN, PTPN11, PTPN13, PTPRB, RAC1, RAD21, RB1, RET, RHOA, RNF43, RPL10, RPL5, RUNX1, SETBP1, SETD2, SF3B1, SH2B3, SMAD4, SMARCA4, SMARCB1, SMO, SOCS1, SPEN, SPOP, SRC, SRSF2, STAG2, STAT3, STAT5B, STK11, SUFU, TBL1XR1, TBX3, TERT, TET2, TNFAIP3, TNFRSF14, TP53, TRAF7, TSC1, TSC2, TSHR, U2AF1, UBR5, USP8, VHL, WT1, XPO1, ZRSR2.

## References

1. Greaves, M. & Maley, C. C. CLONAL EVOLUTION IN CANCER. Nature 481, 306–313 (2012).

2. Yates, L. R. & Campbell, P. J. Evolution of the cancer genome. Nat. Rev. Genet. 13, 795–806 (2012).

3. Andor, N. et al. Pan-cancer analysis of the extent and consequences of intratumor heterogeneity. Nat. Med. 22, 105–113 (2016).

4. Nowell, P. C. The clonal evolution of tumor cell populations. Science 194, 23–28 (1976).

5. Nik-Zainal, S. et al. The Life History of 21 Breast Cancers. Cell 149, 994–1007 (2012).

6. Dentro, S. C., Wedge, D. C. & Van Loo, P. Principles of Reconstructing the Subclonal Architecture of Cancers. Cold Spring Harb. Perspect. Med. 7, (2017).

7. Williams, M. J., Werner, B., Barnes, C. P., Graham, T. A. & Sottoriva, A. Identification of neutral tumor evolution across cancer types. Nat. Genet. 48, 238–244 (2016).

8. Johnson, D. C. et al. Neutral tumor evolution in myeloma is associated with poor prognosis. Blood 130, 1639–1643 (2017).

9. Zack, T. I. et al. Pan-cancer patterns of somatic copy number alteration. Nat. Genet. 45, 1134–1140 (2013).

10. Bozic, I., Gerold, J. M. & Nowak, M. A. Quantifying Clonal and Subclonal Passenger Mutations in Cancer Evolution. PLOS Comput. Biol. 12, e1004731 (2016).

11. Nei, M. & Gojobori, T. Simple methods for estimating the numbers of synonymous and nonsynonymous nucleotide substitutions. Mol. Biol. Evol. 3, 418–426 (1986).

12. Goldman, N. & Yang, Z. A codon-based model of nucleotide substitution for protein-coding DNA sequences. Mol. Biol. Evol. 11, 725–736 (1994).

13. Martincorena, I. et al. Universal Patterns of Selection in Cancer and Somatic Tissues. Cell 171, 1029–1041.e21 (2017).

14. Forbes, S. A. et al. COSMIC: somatic cancer genetics at high-resolution. Nucleic Acids Res. 45, D777–D783 (2017).

15. Williams, M. J. et al. Quantification of subclonal selection in cancer from bulk sequencing data. Nat. Genet. 1 (2018). doi:10.1038/s41588-018-0128-6

16. Hammal, O. A., Alonso, D., Etienne, R. S. & Cornell, S. J. When Can Species Abundance Data Reveal Non-neutrality? PLOS Comput. Biol. 11, e1004134 (2015).

17. Herzog, H. A., Bentley, R. A. & Hahn, M. W. Random drift and large shifts in popularity of dog breeds. Proc. R. Soc. B Biol. Sci. 271, S353–S356 (2004).

18. Leigh, E. G. Neutral theory: a historical perspective. J. Evol. Biol. 20, 2075–2091 (2007).

## References

1. Williams, M. J., Werner, B., Barnes, C. P., Graham, T. A. & Sottoriva, A. Identification of neutral tumor evolution across cancer types. Nat. Genet. 48, 238–244 (2016).

2. Bozic, I., Gerold, J. M. & Nowak, M. A. Quantifying Clonal and Subclonal Passenger Mutations in Cancer Evolution. PLOS Comput. Biol. 12, e1004731 (2016).

3. Van Loo, P. et al. Allele-specific copy number analysis of tumors. Proc. Natl. Acad. Sci. 107, 16910–16915 (2010).

4. Varela, I. et al. Exome sequencing identifies frequent mutation of the SWI/SNF complex gene PBRM1 in renal carcinoma. Nature 469, 539–542 (2011).

5. Jones et al. cgpCaVEManWrapper: Simple Execution of CaVEMan in Order to Detect Somatic Single Nucleotide Variants in NGS Data. Curr. Protoc. Bioinforma. 56, 15.10.1–15.10.18 (2016).

6. Martincorena, I. et al. Universal Patterns of Selection in Cancer and Somatic Tissues. Cell 171, 1029–1041.e21 (2017).

7. Alexandrov, L. B. et al. Mutational signatures associated with tobacco smoking in human cancer. Science 354, 618–622 (2016).

8. Forbes, S. A. et al. COSMIC: somatic cancer genetics at high-resolution. Nucleic Acids Res. 45, D777–D783 (2017).

9. Zoubarev, A. et al. Gemma: a resource for the reuse, sharing and meta-analysis of expression profiling data. Bioinformatics 28, 2272–2273 (2012).

